# Polymer coil-globule phase transition is a universal folding principle of Drosophila epigenetic domains

**DOI:** 10.1101/383158

**Authors:** Antony Lesage, Vincent Dahirel, Jean-Marc Victor, Maria Barbi

## Abstract

**Background:** Localized functional domains within chromosomes, known as *topologically associating domains* (TADs), have been recently highlighted. In *Drosophila*, TADs are biochemically defined by epigenetic marks, this suggesting that the 3D arrangement may be the “missing link” between epigenetics and gene activity. Recent observations (Boettiger et al., Nature 2016) provide access to structural features of these domains with unprecedented resolution thanks to super-resolution experiments. In particular, they give access to the *distribution* of the radii of gyration for domains of different linear length and associated with different transcriptional activity states: active, inactive or repressed. Intriguingly, the observed scaling laws lack consistent interpretation in polymer physics.

**Results:** We develop a new methodology conceived to extract the best information from such super-resolution data by exploiting the whole distribution of gyration radii, and to place these experimental results on a theoretical framework. We show that the experimental data are compatible with the *finite-size* behavior of a *self-attracting polymer*. The same generic polymer model leads to quantitative differences between active, inactive and repressed domains. Active domains behave as pure polymer coils, while inactive and repressed domains both lie at the coil-globule crossover. For the first time, the “*color-specificity*” of both the persistence length and the mean interaction energy are estimated, leading to important differences between epigenetic states.

**Conclusions:** These results point toward a crucial role of criticality to enhance the system responsivity, resulting in both energy transitions and structural rearrangements. We get strong indications that epigenetically induced changes in nucleosome-nucleosome interaction can cause chromatin to shift between different activity states.

## 1 Introduction

Chromosomes are giant polymers [1], *i.e.* very long chains of monomers. In such systems, even very small interactions between monomers can strongly influence the whole structure, as many small interactions can add up to stabilize compact structures. Such polymers can thus exist in more or less compact conformations from swollen coils to collapsed globules, depending on the interaction of the monomers with each other and with the solvent as well as on the temperature. This makes polymer-based (numerical as well as theoretical) modeling approaches more and more popular in describing nuclear architecture.

Chromatin is indeed known to be divided into compartments of various densities, including rather low density regions, generally associated with transcribing genes, and denser ones, more often silent or repressed from the transcription point of view. This spatial compartmentalization is achieved trough linear segmentation of the genome into blocks or *domains*, with a biochemical (*epigenetic*) marking of these domains strongly correlated to their state of activity and spatial folding. Epigenetics, spatial organization and density of the genomic domains must have been tuned by evolution, through the selection of a physical mechanism affecting chromosome folding and transcription activity.

It is not easy, however, to determine the nature and intensity of the underlying interactions in real systems, as well as the presence of specific constraints such as bridges between different chromosomal loci, and their effect. Consequently, a quantitative description well grounded in the molecular level is a crucial issue. Traditional optical imaging techniques cannot be used for this purpose, since their resolution is limited by diffraction to a few hundred nanometers. This limitation has been overcome by the use of super-resolution imaging, as recently achieved notably by Zhuang’s and Nollmann’s groups [2, 3]. Now the question is how to take full advantage of these data in order to catch the underlying physical parameters. Based on a finite-size polymer model, we propose here a theoretical framework enabling to reproduce and interpret super resolution imaging data of chromatin.

To this aim, we first needed to find the good level of description of the functional organization of the nucleus. Topologically associating domains (TADs) are one emerging keystone in the description of the complex and dynamical spatial arrangement of chromosomes, hence gene regulation and cell differentiation. TADs are identified thanks to chromosome conformation capture techniques and may be defined as genomic regions whose DNA sequences physically interact with each other more frequently than with sequences outside the TAD [4]. In *Drosophila*, TADs are equivalently identified by special combinations of biochemical marks called epigenetic states or *colors* [5]. Epigenetic coloring is also specific to different gene activity states [6], this suggesting that the 3D arrangement may be the “missing link” between epigenetics and gene activity. The epigenetic domain level appears then particularly relevant for taking full advantage of 3D measurements. Interestingly, *Drosophila* nuclear organization at the epigenetic domain level has been investigated using SIM [3] or STORM [2]. This allowed in particular to measure the radius of gyration of each individual snapshot for every imaged domain [2]. This provided access to the *distribution* of the radii of gyration for domains of different linear length and associated with different epigenetic states: active, inactive or repressed. As more data will follow this pioneering work, we attend here to define the best methodology to extract information from series of images of equilibrium conformations of polymers.

In recent years *Drosophila* chromosomes have been successfully modeled as block copolymers: each block corresponds to an epigenetic domain and each monomer interacts preferentially with other monomers of the same epigenetic type [7]. The number of DNA base pairs per monomer is arbitrary and depends on the spatial coarse-graining that is chosen. However there is a characteristic lengthscale, the so-called Kuhn length, which measures the polymer intrinsic rigidity: polymer segments smaller than the Kuhn length can be considered as rigid rods. More precisely, in the high temperature/zero interaction limit a real polymer behaves as a freely jointed chain of Kuhn segments. However, in the case of chromatin, the numbers of base pairs contained in a Kuhn segment of length *K*_nm_ (thus expressed in nm) may vary, depending on its linear compaction *c* in bp/nm. Consequently, the Kuhn length expressed in bp is *K*_*bp*_ = *c K*_*nm*_. Then one of the parameters *c* or *K*_*bp*_ should be used in order to compare the model with data expressed in bps. Finally, the interaction parameter −*ε* measures the effective interaction strength *per Kuhn segment* and can be seen as a global parameter accounting for multiple effects, possibly different in different epigenetic states. The three physical parameters, namely the Kuhn length *K*_*nm*_, the linear compaction *c* and the interaction energy −*ε*, completely characterize the epigenetic domain in this theoretical framework.

Strikingly, our approach allows us to determine these three physical parameters from the experimental distributions of gyration radii by a fitting procedure. The main outcome of this comparison is that evolution appears to have selected the folding state of chromosomal domains so to be close to the coil-globule transition, hence to be in the vicinity of the critical point of this second-order phase transition. *Criticality* being a key feature to drive the system in a highly responsive state (characterized by huge susceptibilities), the selected regime is expected to dramatically affect physiological parameters as for example the promoter-enhancer distance, and more generally transcription initiation, by means of small variations of global parameters.

More precisely, our interpretation framework makes it possible to extract from the newly available data estimates of the interaction energy −*ε*, which are remarkably close to coil-globule crossover for the three different epigenetic states (with active domains on the coil side, inactive just at the transition, and repressed on the globule side). Based on our quantitative results, we can draw inferences that *nucleosome-nucleosome* tail-bridging interaction is most likely at the origin of the system’s criticality. This result is clearly consistent with previous observations in solutions of isolated nucleosome core particles [8, 9]. It suggests that the evolution of chromatin folding might have been driven by a very fine-tuned optimisation of nucleosome-nucleosome interactions, so as to be close to the coil-globule crossover.

## 2 METHODS

### 2.1 Theoretical framework of polymer physics

We model an epigenetic domain a as polymer chain made of *N* identical monomers, of position *r*_*i*_, interacting by contacts with their nearest neighbours (see Fig. 1). A standard way to quantify the mean size of a single-polymer chain in a given configuration is the so-called radius of gyration *R*_*g*_, defined as the standard deviation of its position the distribution of its monomer positions:

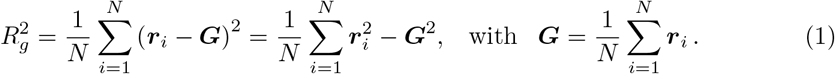

It is common to statistically characterize the average behavior of a polymer of *N* monomers by means of the mean radius of gyration,

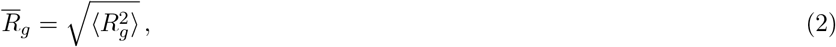

where the average 〈·〉 is performed over the ensemble of conformations for a given polymer. The *scaling behavior* of 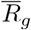 with the polymer length *N* reads

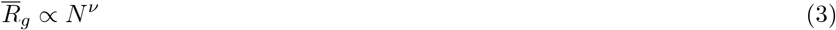

where the scaling exponent *ν* is the so-called *Flory exponent*.

For a polymer at equilibrium and in the large-*N* limit, two different folding modes have been predicted and measured [10], depending on the relative strength of the interaction energy with respect to temperature *ε/k*_B_*T*: In good solvent (low *ε/k*_B_*T*), the favorable interaction with the solvent leads to an effective repulsion between monomers. Hence, the polymer expands into a decondensed, disordered state called *coil*, described as a *self-avoiding walk* (SAW); In poor solvent (high *ε/k*_B_*T*), monomer-monomer attractions become predominant, and the polymer collapses into a state called *globule*. The phase transition between the two regimes is observed at a specific temperature called Θ (*theta*) temperature or Θ *point*, or equivalently at *ε*_θ_ = *k*_*B*_Θ. At this point, the effective repulsion between monomers compensates their attraction [11, 12]. The polymer behaves then as a random walk (RW) (also referred to as Θ-polymer). Each of the three regimes is characterized by a specific Flory exponent: the three values are summarized in Table 1.

**Table 1.** Summary of conditions and Flory exponents expected for the three typical polymer folding states.

|  | Coil (SAW) | Theta-polymer (RW) | Globule |
| --- | --- | --- | --- |
| Condition: | $\varepsilon < \varepsilon_\theta$ | $\varepsilon_\theta = k_B \Theta \simeq 0.27$ | $\varepsilon_\theta < \varepsilon$ |
| Flory $\nu$: | 3/5 | 1/2 | 1/3 |

### 2.2 Finite-size polymers display a richer scaling behavior

Self-attracting polymers of *finite-size N* undergo a coil-globule transition at a *N-dependent* critical temperature Θ_*N*_ < Θ (or critical energy 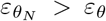) [13, 14]. Moreover, the abrupt phase transition is replaced by a *crossover*, i.e. a gradual variation of the mean radius of gyration 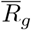 as a function of *ε* at fixed *N*. The scaling behavior of 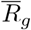 as a function of *N* at a given *ε* is, consequently, also affected (see Fig. 1).

Using a refined version of the semi-empirical *finite-size polymer theory* first introduced by one of us [15, 16], we were able to express the polymer free energy 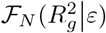 as a function of its instant radius of gyration [17]. The resulting expres ion for the free energy is more easily expressed in terms of the renormalized density *t* = *ρ*^1/(*νd*−1)^ where 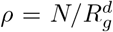 is the local monomer density and *d* = 3 the space dimension. The free energy reads then

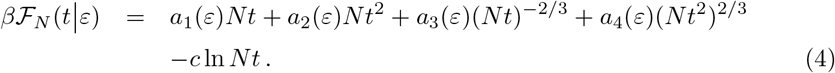

Most importantly this formula factorizes the *N*-dependence (i.e. the finite-size effects) and the energy-dependence. It is designed as a renormalized Flory free energy. The term *a*_1_(*ε*) plays the role of a second virial coefficient, vanishing at the Θ *point* (i.e. for *ε* = *ε*_θ_). The second term *a*_2_(*ε*)*Nt*^2^ accounts for three-body interactions which become dominant in the globule phase. The third one *a*_3_(*ε*)(*Nt*)^*−*2/3^ is relevant for extended conformations and the term *a*_4_(*ε*)(*Nt*^2^)^2/3^ accounts for surface tension effects which become crucial in the coil-globule crossover region. The logarithmic correction −*c* ln*Nt* is related to the so-called enhancement factor of self-avoiding walks, with *c* = −1.13 [18].

Having the free energy as a function of the gyration radius makes it possible to obtain the *distribution* of this quantity, for different *N* and *ε*, namely 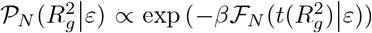. The four coefficients *a*_1_(*ε*)…, *a*_4_(*ε*) have been fitted on the distributions of gyration radii obtained from simulations at different *N* and *ε* (supplementary Fig. S2) [17]. The corresponding, theoretical *average* values 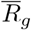 and typical configurations are reproduced in Fig. 1 ^[1]^.

The most striking feature of Fig. 1 is the slope inflection when *ε* > *ε*_*θ*_. In this regime, 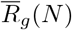 displays a characteristic knee-point around some value *N* = *f* (*ε*/*k*_*B*_*T*) [13] which defines the crossover region. In this region, the radius of gyration hardly changes, giving *e.g.* close radii of gyration for the *N* = 109 (coil) and *N* = 538 (globule) blue snapshots. This behavior is quite unusual among critical phenomena and leads to dramatic finite-size effects. Remarkably, the same kind of behavior is observed in the case of a block copolymer, where block conformations are affected by finite-size effects likewise isolated polymers [20]. Similar effects can thus be expected in the case of epigenetic domains embedded in larger chromosomal regions. Fig. 1 also clarifies the difference between the finite-size and large-*N* descriptions: for sufficiently large *N*, indeed, only two scaling regimes remain, corresponding to coil (resp. globule) conformations if *ε* < *ε*_*θ*_(resp. *ε* > *ε*_*θ*_).

### 2.3 Mapping experimental data on the adimensional theoretical model

The aforementioned model relates dimensionless quantities, namely the number of monomers *N* and the radius of gyration *R*_g_ expressed in monomer, or Kuhn length, units. In the case of chromosome domains, the number of monomers is unknown whereas the measurable physical parameters are the domain spatial extent 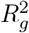 measured in nanometer and the linear length *L* of the domain measured in base pairs. A mapping of the dimensional data on the adimensional model is then a crucial step to the aim of using the theoretical model to infer physical parameters from experiments. The two Kuhn lengths *K*_nm_ and *K*_bp_play the role of rescaling parameters in this mapping: *K*_bp_ relates the number of monomers *N* to *L* by *N* = *L/K*_bp_, and *K*_nm_ provides a physical length scale to the size distribution predicted by the model.

**Figure 1.**
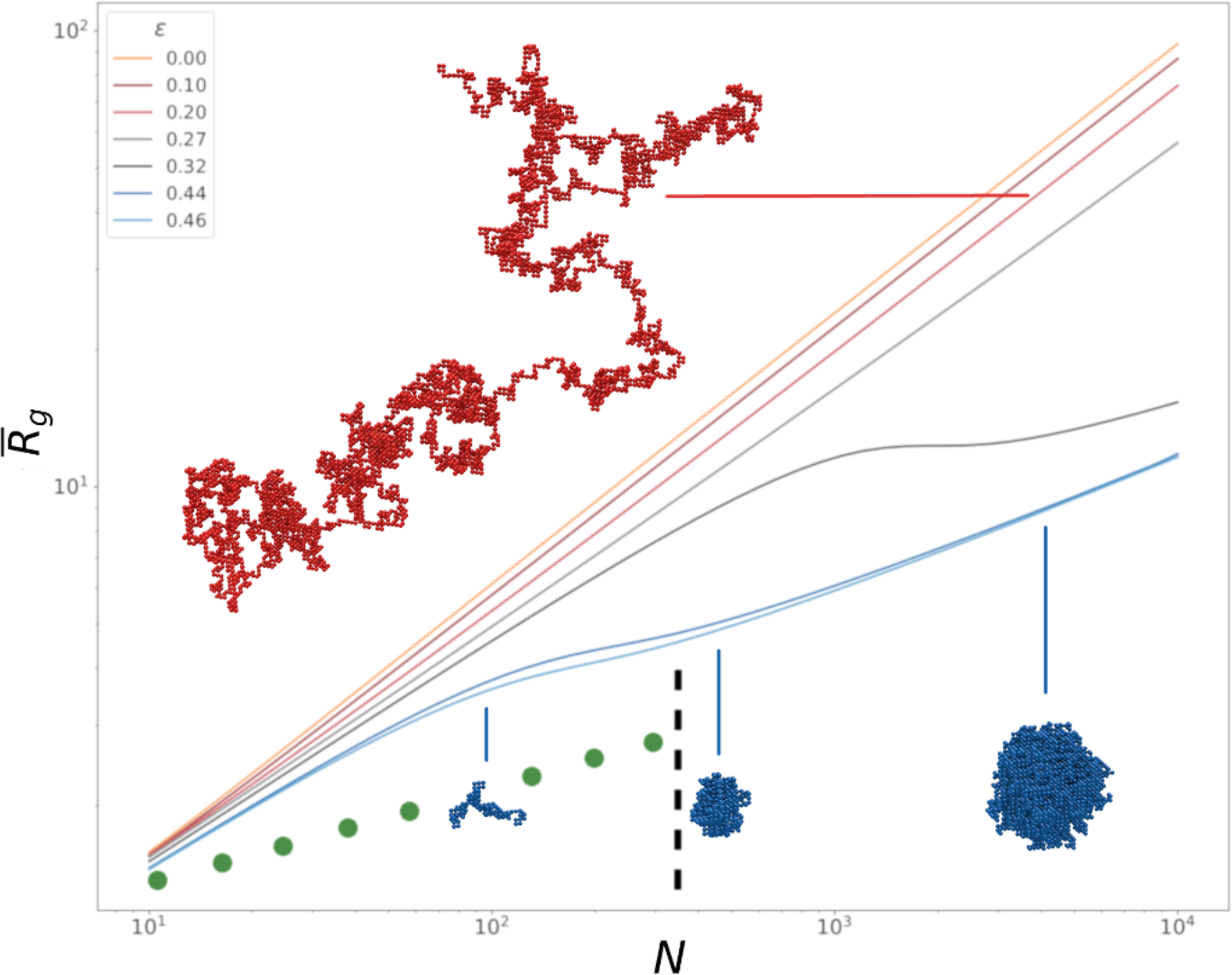
Theoretical curves for a finite-size polymer model. Log-log plot of mean radii of gyration 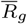 (2) (in Kuhn length units) against the number of monomers *N* at different values of *ε* below and above *ε*_θ_ ≃ 0. 27. Typical configurations at *ε* = 0. 20 *k*_*B*_*T* and *N* = 5012 (red), *ε* = 0. 44 *k*_*B*_*T* and *N* =109, 538 and 5012 (from left to right, blue) are shown. The green dots are a rough representation of the radius of gyration as a function of the domain length from the simulated sticky polymer domain of Ref. [2] (Fig. 4.c), up to the close packing limit (black dashed line).

As already mentioned in the Introduction, the correspondence between *K*_nm_ and *K*_bp_ is *a priori* not known. It depends on the local chromatin linear compaction in bp/nm *c*, as *K*_bp_= *c K*_nm_, and can thus vary in different domains. The compaction *c* is difficult to estimate, because the nucleosome fiber architecture is not directly observable. We then considered *K*_nm_ and *K*_bp_ as two independent parameters of our model, in addition to *ε*, the interaction energy between Kuhn segments. We relied, however, on the plausible hypothesis that *c* is homogeneous *within* one epigenetic domain and is the same for all domains of the same *color*.

In practice, the mapping on dimensional measurements results in a reformulation of the free energy in order to use *K*_bp_ and *K*_nm_ explicitly as fitting parameters. It yields the following expression for the probability density function of 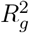

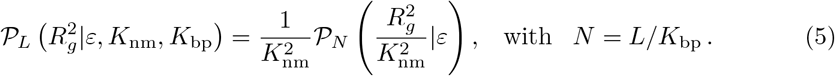

### 2.4 Dataset and statistical analysis

Boettinger and co-workers provided us with the ensemble of their radius of gyration measurements. These authors identified candidate domains of a specific length *L* by applying a moving average filter with a window of same size *L* on the marker enrichment trace for the marker of the desired epigenetic state. The whole dataset consists in three sets of data for the three different epigenetic *colors*: active (*red*), inactive (*black*) and repressed (*blue*). These three data sets contain 23, 14 and 11 domains of different lengths, respectively. For each of the 48 domains, the radius of gyration is measured over a set of 20-100 cells.

For a given *color*, we assume a unique set of parameters *θ* is needed. Then, we define the log-likelihood of the measured set of radii of gyration 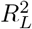 for the given length *L*,

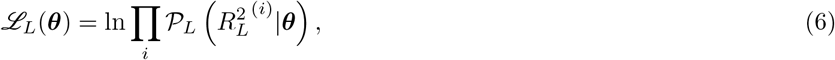

where 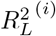 is the *i*-th measured radius of gyration. Finally, the *total* log-likelihood *ℒ* = ∑_*L*_ *ℒ*_*L*_ allows to infer the distribution of the parameters *θ*, over all lengths of a given *color*, with the Goodman & Weare’s Affine Invariant Markov chain Monte Carlo Ensemble sampler [21]. We used as the best estimate of the parameters their mathematical expectation according to the distribution previously obtained and confidence intervals have been deduced, as well, by evaluating their standard deviation. Parameters positiveness has been insured by a uniform prior distribution.

## 3 RESULTS

### 3.1 Power law fit of experimental gyration radii leads to unusual exponents

We used the full ensemble of measurements of Ref. [2] to analyze the scaling of the *mean* and *median* radii of gyration for *Drosophila* domains of different lengths and belonging to three epigenetic states: (i) active *red* types, covering the expressed regions, (ii) inactive *black* states and (iii) repressed *blue* domains, characterized by the presence of Polycomb group (PcG) proteins. The ensemble of resulting power law exponents are given in Table 2. Corresponding plots are given in supplemental Fig. S1. Surprisingly enough, the inactive and repressed datasets display scaling exponents *ν* of 0. 30 and 0. 21, which are both smaller than the expected value of the globular state *ν* = 0. 33, while the active dataset is fitted with *ν* = 0. 34. Also intriguingly, the apparent *ν* exponents of the median for the three domain types are larger than for the mean (*ν* = 0.37, 0.30, 0.24 for active, inactive and repressed, respectively). For large-*N* polymer conformations, the scaling is expected to be identical for the median and the mean. We thus obtain here a rather strong indication of a finite-size effect.

**Table 2.** Summary of scaling exponents obtained from a power law fit of data from Ref. [2] for active (A), inactive (I) and repressed (R) epigenetic domains, of either the mean or median values of the radii of gyration for all different *colors* and lengths.

| State: | Active | Inactive | Repressed |
| --- | --- | --- | --- |
| Means | $0.34 \pm 0.02$ | $0.30 \pm 0.02$ | $0.21 \pm 0.01$ |
| Medians | $0.37 \pm 0.02$ | $0.30 \pm 0.03$ | $0.24 \pm 0.02$ |

These unexpected results are partially accounted for in Ref. [2] by means of simulations where inactive and repressed domains are modeled with a mixing of *sticky* and *non sticky* polymers prepared in special initial conditions and let relaxed in a confined volume. If this approach makes it possible to reproduce the *exponents* observed for the three types of domains respectively (at least for the medians), it does not make it possible to reproduce the observed values of 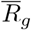 *quantitatively*. Furthermore, the repressed domains occur to be almost close packed, inactive domains compaction is essentially due to imposed confinement (at zero attractive interaction energy), and the difference between active and inactive domains is not taken into account. We therefore decided to address the question of how to explain these unexpected findings.

### 3.2 Bundle correction for tetraploidy is necessary but not sufficient

A possible explanation for the observed anomalies may arise from the particularity of the cell line used in these experiments. The chromosomes of the tetraploid Drosophila Kc167 line are known to form bundles, sticking together in a regular fashion with a pairing rate of about 80% [22, 23]. Super-resolution imaging techniques do not distinguish paired chromosomes without allele-specific labeling (this is however in principle possible by STORM experiments for small enough domains, and has been done by SIM [24]).

In order to take tetraploidy into account, we propose to describe domains as bundles of *n* Kuhn chains of *N* segments. The resulting radius of gyration reads

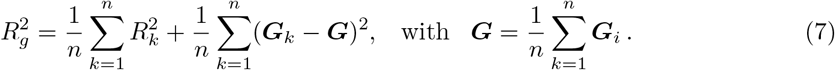

where *G*_*k*_ is the center of mass of the *k*-th polymer of the bundle and 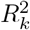 its radius of gyration. The second sum in Equation 7 is the bundle contribution to the total radius of gyration, 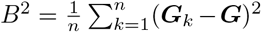. Taking the average of Equation 7 leads to 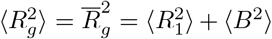 Hence, the bundle effect expected on 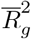 is essentiallya constant positive shift.

We inferred *B*^2^ from the experimental distribution obtained for the smallest epigenetic domains, whose mean radius of gyration is mainly determined by the bundle extension. Including this correction to the standard power laws described above will, by construction, resolve the observed discrepancies for domains of small length, but it is not sufficient to fix the whole ensemble of observations (data not shown). It therefore seems necessary to introduce a more accurate modeling of the system.

### 3.3 All the 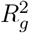 distributions are fitted by a unique finite-size polymer model

In dealing with the difficulties just enlightened, two remarks could be made as a matter of principle. First, previous analyses take into account the mean (or median) of the measured quantities. However, a full distribution of measurements for each domain length and epigenetic state is available: this large dataset should be fully analysed to achieve a better interpretation of the experimental results. Second, the modelling proposed so far has only considered the polymer behavior in the large-*N* limit. Finite-size polymers display, however, a much richer behaviour, that may potentially introduce new interesting features for the comparison with data. We therefore decided to take into account both features at a time by adopting a finite-size polymer model and looking for the whole *distribution of the radius of gyration*.

The correction for tetraploidy previously introduced for the *mean* radius of gyration should be then extended to the *distribution*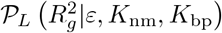. To do this, we inferred the whole distribution of the bundle size *B*_2_ from the distribution observed for the shortest domains, that is dominated by the bundle contribution. For this domains, the distribution displays a shifted exponential form

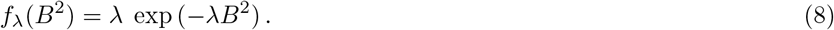

This is compatible with a random arrangement of the four polymers, with a dispersion *σ*^2^ = 1*/λ* characteristic of the bundle section spreading. The bundle section spreading may depend in general on the polymer length: *σ*^2^ = *σ*^2^ (*N*). We chose to model *σ*(*N*) as varying from a minimum value *a*_0_to a maximum value *a*_∞_, reached within a characteristic length scale *N*_0_:

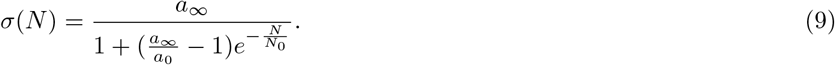

We rely on the approximation that the bundle extent *B*^2^ and the one-polymer radius of gyration 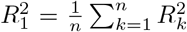 are independent random variables. Hence, we finally write the observed distribution as the convolution of the theoretical, single-polymer distribution and of the bundle function:

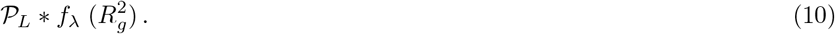

To analyze the data, we performed a Markov Chain Monte Carlo (MCMC) Bayesian inference (Section 2.4), given the the observation data, in order to infer the parameters of our theoretical finite-size self-avoiding polymer (*ε*, *K*_*bp*_, *K*_nm_) (Eq. (5)), corrected for tetraploidy with the bundle parameters (*a*_0_, *a*_∞_, *N*_0_). Importantly, we rely on the following assumptions:

i. the *same* general polymer model can describe all the observations whatever the epigenetic state, or *color*
ii. different *colors* correspond to different sets of model parameters;
iii. the ensemble of domains of a given *color* can be fitted with a unique set of parameters, whatever the size of the domain, its genomic context, or other characteristics.

Datasets for each of the three epigenetic *colors* are therefore analyzed as a whole.

Probability distributions for the six parameters and for the case of active, inactive and repressed domains are displayed in supplemental Fig.s S3, S4 and S5, respectively. The joint posterior distributions for any pair of parameters show in particular the absence of correlation between the bundle parameters and the energy parameter *ε* in these distributions.

In Fig. 2, all the fitted histograms are plotted along with the theoretical curves obtained with the optimal parameters. Separate histograms are given in Fig.s S6, S7 and S8 for the three *colors* respectively. Compatibly with the limited size of the experimental dataset, the comparison shows a remarkably good agreement between the distribution of data and the predicted behavior.

**Figure 2.**
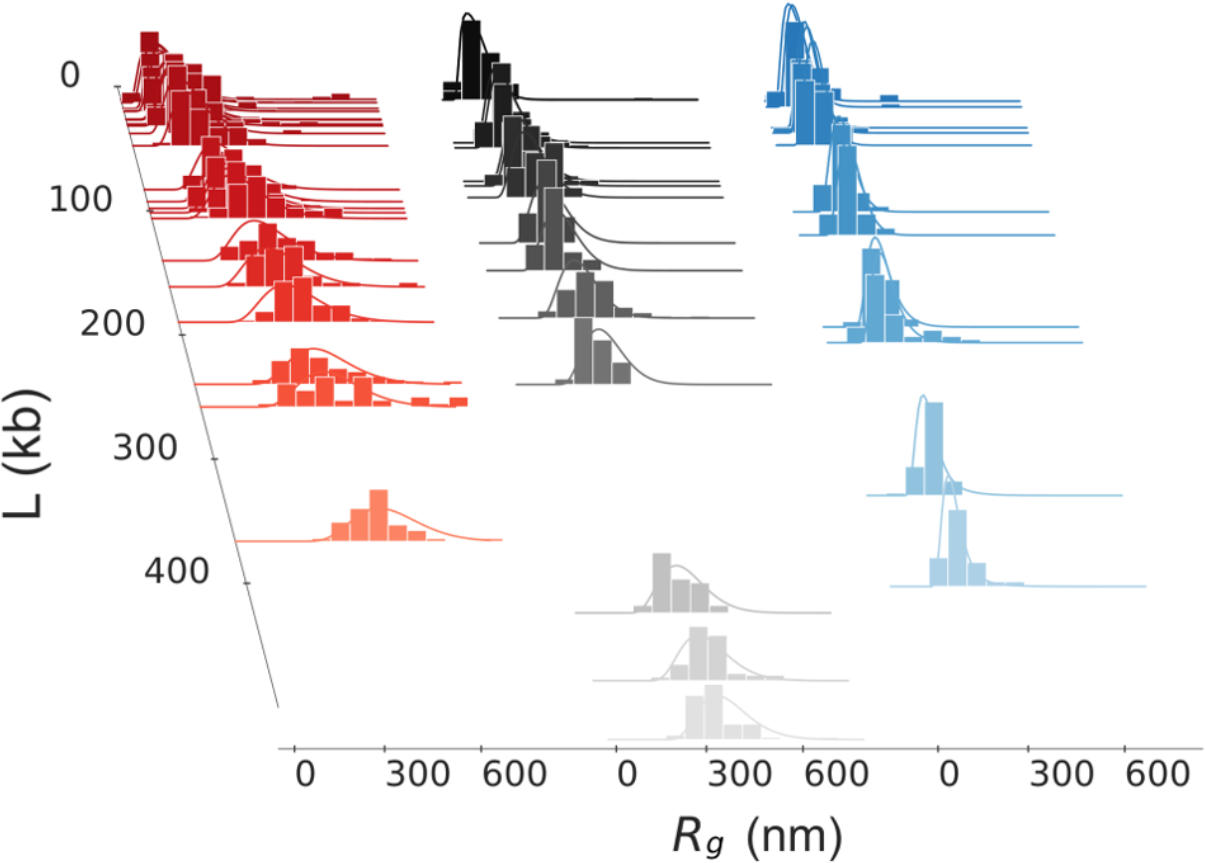
Experimental data fit: distributions. The three data ensembles from Ref. [2] (histograms) with the corresponding theoretical fitting distributions (lines). *Colors* refer to epigenetics: red for active, black for inactive and blue for repressed domains. The theoretical distributions have been calculated from the analytical expression of the probability density by using the fitting parameters of Table 3. A more detailed view of the complete set of histograms and fits is given in Fig.s S6, S7 and S8.

The fit of the gyration radius distributions allows the data to be used in the most complete way and allows the maximum amount of information to be extracted. As an *a posteriori* check of the results, the theoretical *mean* radius of gyration 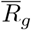 as a function of the domain length *L* can be compared to the experimental averages. This is done in Fig. 3**A**.

**Figure 3.**
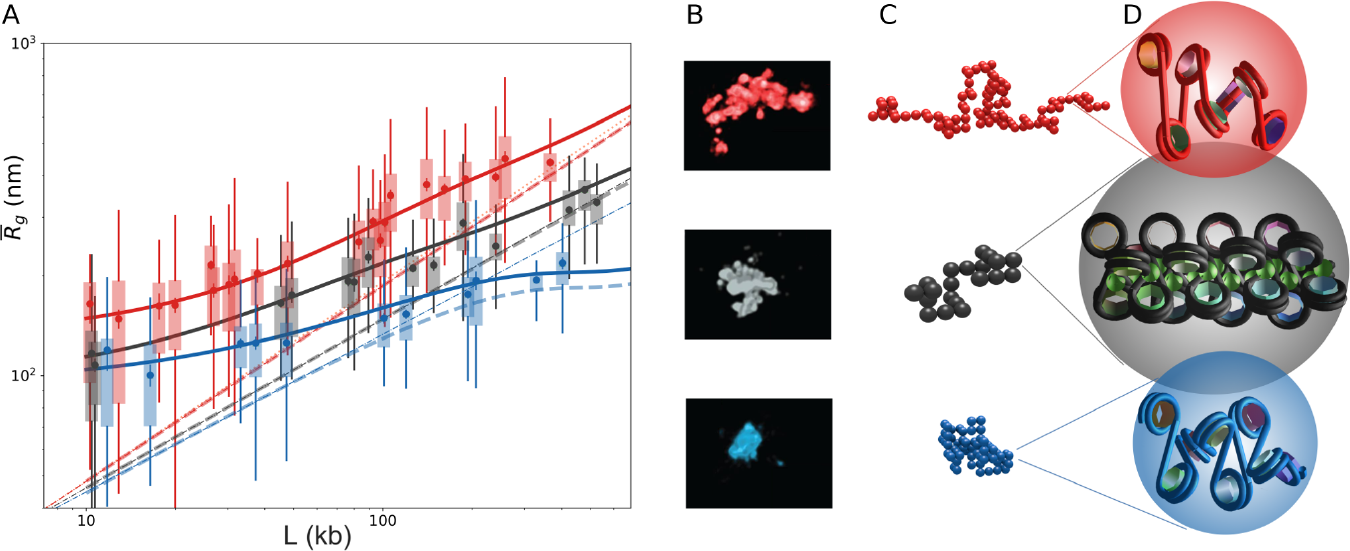
A. Experimental data fit: mean gyration radii. Mean 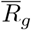 as a function of the domain length *L* calculated from the analytical model with the parameter sets of Table 3: active (red line), inactive (black line), repressed (blue line). Boxplots (same *colors*) correspond to the experimental data from Ref. [2]. Dashed lines are obtained from previous fitting curves by deconvolution, hence correspond to the behavior expected in an haploid system. The orange dotted line represents the coil (*ν* = 3/5) typical scaling law. A corresponding fit for *median* values is given in Supplemental Fig. S9; **B. Experimental images.** 3D-STORM images adapted from Boettiger *et al.* [2] corresponding to an active, inactive and repressed domain (from top to bottom; 106, 79 and 119 kb respectively). **C. Fitting model snapshots.** Typical configurations of the domains shown in (B) obtained with the corresponding fitted parameters *K*_nm_, *K*_bp_and *ε*; **D. Corresponding monomers at the fiber scale.** Twoangle models of the nucleosome fibers corresponding to the fitted parameters of the domains shown in (B) and simulated in (C). In the case of *black* domains, the green spheres suggest the presence of H1 histones.

Interestingly, we could here get rid of the bundle on the fitting curves by applying a deconvolution procedure, thus predicting what would be observed with a haploid genome. The resulting curves are shown in Fig. 3**A** as dashed lines. Overall, these results show that our method, together with the assumptions made, gives a correct fit of the whole dataset, and allows for a physically sound interpretation which will be commented in the following.

### 3.4 Attraction brings inactive and repressed domains close to critical conditions

Fig. 3**A** clearly shows that active domains have a scaling exponent very close to 3/5 (*red dashed lines*) and stay thus in the coil regime for all the observed lengths. This is in agreement with the fitted parameter *ε* = 0. 10 *k*_*B*_*T* obtained for active domains, well below the theoretical (large-*N*) transition value of *ε*_*θ*_ ≃ 0. 27 *k*_*B*_*T*.

Table 3 summarizes the parameters obtained for the three epigenetic states, together with the derived linear compaction in different units. At variance with active domains, the repressed (*blue dashed lines*) domains are above the critical energy, with *ε* = 0. 44 *k*_*B*_*T*, and are theoretically in the globule side of the transition. However, the *finite-size* critical energy is larger than its large-*N* limit *ε*_*θ*_: repressed domains are in fact in the crossover region. In Fig. 3**A**, a plateau is indeed visible around lengths of ~400 kb, with a clear-cut crossover from coil to globule behavior.

As somehow expected, inactive (*black dashed lines*) domains display an intermediate regime: With *ε* = 0. 32 *k*_*B*_*T*, they are above the limit coil-globule transition energy but finite-size effects remain strong at the observed lengths. As a result, the crossover plateau is still not reached at these lengths, but a clear discrepancy with respect to the short-range coil behavior is observed.

In any case, all short domains are close to coil conformations due to finite-size effects, that emerge then as a crucial feature in the interpretation of domain super-resolution imaging. A fit of the deconvolved curves slopes in the small domain region (< 60 kb) gives indeed *ν* = 0. 51 and 0. 47 for inactive and repressed domains, respectively, *i.e.* a behavior close to the *ν* = 0. 5 RW expected at the transition (we obtain *ν* = 0. 59 for the active deconvolved fit within the same range).

### 3.5 Fitted architecture parameters are compatible with structural chromatin models

Both values of *K*_nm_and *K*_bp_ are obtained simultaneously by our approach. The possibility to determine these structural parameters is a remarkable consequence of the coil-globule crossover. This comes from the existence of different asymptotic scaling laws with *N* before, during and after the crossover. This means, on the other hand, that these parameters can only be obtained if the data allow to explore the crossover region. Therefore, the data for the active domains, which are all below the transition, do not allow these two structural parameters to be determined independently.

For repressed domains, we obtain *K*_nm_ ~ 35 nm and *K*_bp_ ~ 1500 bp (Table 3). The corresponding compaction *c* ~ 40 bp/nm corresponds to *c*_10_ ~ 2 nucl./10 nm. The mean linear compaction of the nucleosome fiber is in principle determined by the underlying architecture of the nucleosome fiber, which in turn depends on a few local parameters, namely the nucleosome repeat length (NRL) and the degree of DNA wrapping around the nucleosome. A simple estimation of the elastic properties and of the compaction of this assembly can be obtained by the two-angle model [26] (Fig. 4). The mechanical and structural features estimated here for repressed chromatin features fit easily with what is analytically obtained in the framework of the two-angle model with standard NRL (192 bp) and crystallographic wrapping angle (negatively crossed nucleosomes). Fig. 3**D** shows, in blue, what the supramolecular architecture of a repressed Kuhn segment looks like, when simply sketched with the two-angle model.

**Table 3.** Summary of physical parameters obtained from the fit of Boettiger’s data [2] for active (A), inactive (I) and repressed (R) epigenetic domains through the Bayesian procedure (mean values, see Fig.s S3 to S5). Errors are calculated from the standard deviations of marginalized parameter distributions. At the bottom, some derived geometrical parameters as the compaction in bp/nm, in nucleosomes per 10 nm, the number of nucleosomes per Kuhn segment *C*. Derived parameters are calculated by assuming a nucleosome repeat length of 182 bp for active domains, 192 bp for inactive and repressed domains [25] (The numerical results obtained with 182 or 192 bp are very close, in the error range). For active domains, the right column estimates are obtained by including architectural features, see Discussion.

| State | Active |  | Inactive | Repressed |
| --- | --- | --- | --- | --- |
|  | Bayesian fit | Estimate | Bayesian fit | Bayesian fit |
| *Fitted:* |  |  |  |  |
| $\varepsilon [k_B T]$ | 0.1 ± 0.05 | | 0.32 ± 0.03 | 0.44 ± 0.04 |
| $K_{bp}$ [bp] | $K_{nm} \propto K_{bp}^{0.56}$ | $\sim 1100 - 1500$ | 3900 ± 1300 | 1500 ± 550 |
| $K_{nm}$ [nm] | | $\sim 32 - 37$ | 60 ± 9 | 37 ± 6 |
| $a_0$ [nm] | 130 ± 7 | | 93 ± 10 | 94 ± 4 |
| $a_\infty$ [nm] | 290 ± 15 | | 170 ± 10 | n/a |
| $N_0$ | 630 ± 370 | | 10 ± 6 | n/a |
| *Derived:* |  |  |  |  |
| $c = \frac{K_{bp}}{K_{nm}}$ [bp/nm] | | $\sim 35 - 40$ | 66 ± 24 | 40 ± 16 |
| $c_{10}$ [nucl./10 nm] | | $\sim 1.9 - 2.2$ | 3.5 ± 1.5 | 2 ± 1 |
| $C$ [nucl./$K_{nm}$] | | $\sim 6 - 8$ | 20 ± 7 | 8 ± 3 |

At variance with repressed domains, inactive domains give *K*_nm_ ≃ 60 nm and *K*_bp_ ≃ 4000 bp, with a corresponding *c*_10_ = 3. 5 nucl./10 nm, hence a nucleosome fiber almost twice as stiff and twice as compact as for repressed domains. In the framework of the two-angle model, such values can only be obtained with an abnormally short NRL, whatever the wrapping. The question then arise of how to justify these findings on a molecular basis. As it will be discussed in 4.2, one possible explanation is given by the stiffening effect of the linker histone H1.

**Figure 4.**
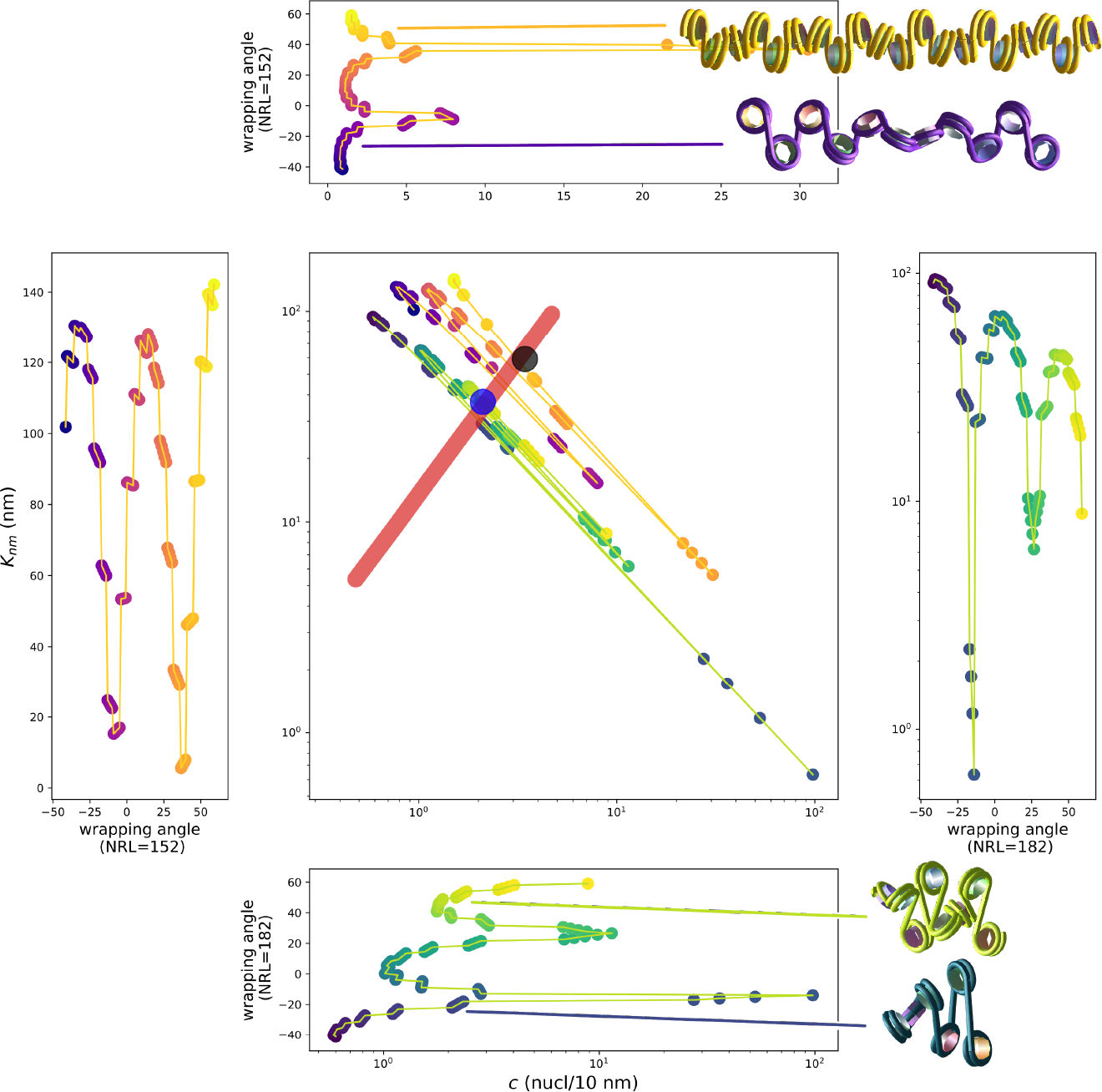
Comparison between the observed behavior and the calculations on a two-angle model of the nucleosome fiber. In the central plot, the Kuhn length *K*n*m* is reported as a function of the compaction *c*10 in nucleosomes per 10 nm. Active domains are represented by the observed power law (red thick line) while for inactive and repressed domains we directly reported the measured values from Table 1 (black and blue dots, respectively). The colored dot-line plots correspond to the same physical quantities calculated on the basis of the two-angle model in a large range of wrapping angles, going from a completely wrapped nucleosome (angle ~ 50°) to a largely open one (angle ~ −30°). The angle dependence is explicitly accounted for in the four lateral plots. Two cases with NRL = 182 bp (blue-green) and NRL = 152 bp (purple-orange) are reported here. The four displayed fibers are completely regular fibers obtained by using the two-angle model for the following parameter sets: angle = 50°, NRL = 152 bp (yellow-orange fiber); angle = −22°, NRL = 152 bp (purple fiber); angle = 50°, NRL = 182 bp (yellow-green fiber); angle = −22°, NRL = 182 bp (blue-green fiber). For each fiber picture, the length corresponds precisely to one Kuhn length as calculated from the corresponding parameter set.

Finally, active domains are in the scale invariant regime (*ε* = 0. 1 *k*_*B*_*T*) where *K*_nm_ and *K*_bp_ cannot be computed independently. They satisfy, instead, such a relation as 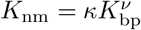 with *κ* some constant and *ν* the Flory exponent. Hence, one would expect the log-likelihood in the (*K*_nm_*, K*_bp_) space to be nearly constant along the curve of equation 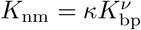. We found indeed a power law fit of the marginalized (*K*_nm_*, K*_bp_) distribution of the form 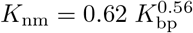 with an exponent very close to the expected Flory exponent (Fig. S3). The log-likelihood in the (*K*_nm_*, K*_bp_) plane is indeed nearly constant along this power-law curve. As a consequence, both averages *K*_nm_and *K*_bp_ obtained from the marginalized distributions for active domains are ill-defined, and they are indeed unrealistically small. In order to identify reasonable ranges for both *K*_nm_and *K*_bp_in active domains, we resorted once again to the two-angle model to calculate the geometry-based 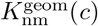 as a function of *c* for any given NRL and wrapping angles. We then replaced *K*_bp_ = *cK*_nm_ in 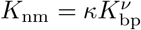 so to obtain *K*_nm_(*c*) from the data. An intercept between 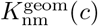 and *K*_nm_(*c*) exists for relatively open wrapping angles, typical of *open* nucleosomes, and with the expected NRL of 182 bp [25] (Fig. 4). The intercept gives *K*_nm_ ~ 35 nm and *K*_bp_ ~ 1300 bp (Table 3). The corresponding compaction *c* ~ 35 bp/nm corresponds to *c*_10_ ~ 2 nucl./10 nm (Fig. 3**D**, red fiber). Interestingly and rather surprisingly, we found by this procedure that the geometrical parameters of active domains are essentially indistinguishable from what previously derived for repressed domains. If confirmed, this finding seems to indicate that active and repressed domains are in fact very close from a structural point of view, and differ essentially only with respect to the interaction energy *ε*.

To sum up these findings, Figures 3**B** and **C** compare typical STORM images [2] with typical configurations at the corresponding parameters, *i.e.* by using the parameters of Table 3 and a number *N* of monomers corresponding to the length of the images domain. Figure 3**D** reproduces the corresponding monomer stretch as obtained with the two-angle model, showing at a glance its physical size and linear density.

To end, it may be useful to point out here that the regular and somehow rigid nucleosome arrays of Figures 3**D** and 4 should only be intended as indicative representations of *average* organizations: the actual nucleosome orientation depends of course on precise architectural parameters that display inhomogeneities along the genome [27, 28], and are, moreover, dynamically fluctuating [29–31]. Note also that the two-angle model is used here only to check the plausibility of physical parameters that are obtained in a completely independent way by means of a much more coarse grained model, i.e. the interacting polymer model.

### 3.6 Bundle geometry

The fitting parameters accounting for the bundle geometry are independent from Kuhn lengths and energy parameters, as shown by the marginal distributions (see Fig.s S3, S4 and S5). As shown in Table 3, we obtain minimal bundle section extents of the order of 100 nm for the three *colors* (with a slightly larger value for active domains) compatibly with the similar radii of gyration observed for small domains (Fig. 3**A**). Interestingly, however, the variation of the bundle section as a function of domain lengths significantly differs for different epigenetic *colors*. Active domains appear to allow for the largest bundle section spreading, up to *a*_∞_ ~ 300 nm provided that the polymer is long enough (*N*_0_ being of the order of 500 monomers, i.e. approximately 15000 nm or 600 kb). In the case of repressed domains, the bundle section spreading approach results in diverging values of *N*_0_ and *a*_∞_, indicating that the simplified constant section description *σ*(*N*) = *a*_0_ is more consistent with the data. We therefore used this simpler model in this case to get a good convergence for the remaining parameters. These findings reveal a correlation between the bundle geometry and the folding mode of the domain. This seems logical since both relate to the extension of the polymer. In the extreme case of a condensed globule, we do not expect that the density inside the domain fluctuates: it is homogenous, of constant density. As a consequence, a bundle of several chains is also uniformly packed and its extent is not related to the polymer length. On the opposite, in coils, chains do not stick together and the interchain distance fluctuates. These fluctuations rise with the length of the chain. This explains why the rather decondensed active domains spread more for longer chains, while this spreading is less intense for more globular inactive and repressed domains.

## 4 DISCUSSION

### 4.1 Obtained parameters support short persistence lengths

Estimates of Kuhn lengths in different organisms remain elusive, one of the major obstacles lies in the need to relate the Kuhn lengths in nanometers and basepairs, through the linear compaction of the chromosome. Few *in vivo* measurements of Kuhn lengths have been made up to now. In 2004, Bystricky *et al.* obtained by high-resolution imaging techniques Kuhn lengths of the order of 400 nm and linear densities of 6–9 nucleosomes per 10 nm in budding yeast [32]. In 2008 Dekker obtained, by 3C and again in yeast, Kuhn lengths of the order of 120–260 nm and linear densities of about 1.1–2.2 nucleosomes per 10 nm [33]. Importantly, the Kuhn length strongly depends on the linear density: fibers of such compaction, the Kuhn length never exceeds 100 nm [26], which is the lower bound of Dekker’s measurements. Moreover, smaller linkers result in less flexibility [26]. Linkers are smaller in yeast than in *Drosophila* (and human) we expect therefore Dekker’s estimates to be an upper bound of the Kuhn length in *Drosophila*. Recent images of chromatin in vivo also confirm low mass densities and highly flexible nucleosome fibers [27].

More recently, Hi-C cyclization and contact probability *P* (*s*) outcomes in mammals suggest a Kuhn length in base pairs of roughly 1 kb for chromatin fibers and certainly less than 5 kb, this suggesting that at the scale of the typical gene (~15 kb), chromatin is highly flexible [34]. This flexibility is also compatible with (and essential for) loop formation via extrusion. In *Drosophila*, same results are obtained by Hi-C high-resolution measurements (G. Cavalli, personal communication).

Taken together, results are thus compatible with linear densities of the order of 2 nucleosomes per 10 nm, and Kuhn lengths as low as *K*_nm_~ 30 nm and *K*_bp_~ 1 kb, in agreement with our results. The relatively small values of *K*_nm_(30 to 60 nm), as compared with naked DNA in particular, confirm the most recent dynamic measurements of the high flexibility of chromatin in vivo (Socol *et al.*, bioRxiv 192765). Note that similar estimates are also used in recent modeling works including in the modeling part of Boettiger’s paper, although not all of them at once [2, 35].

### 4.2 Parameter dependence on epigenetic *colors* points toward a special structure for inactive domains

In previous studies, and notably in simulations of 3D genome organization, it has generally been assumed an unique size of the monomer (*K*_bp_ or *K*_nm_) whatever the epigenetic state. One of the important findings of our work is that active (*red*) and repressed (*blue*) domains have indeed, though surprisingly, the same monomer size (*K*_nm_~ 35 nm and *K*_bp_~ 1500 bp), whereas inactive (*black*) chromatin has a monomer size (*K*_bp_ or *K*_nm_) about twice as large.

As *blue* chromatin domains are dispersed among the volume of the so-called active compartment [3], the nucleosome fiber structural similarity of active and repressed domains may facilitate transitions between active and repressed epigenetic states in the course of cell differentiation.

The increased compactness and stiffness found for inactive domains needs for a justification on a molecular basis and requires a more in-depth discussion. Interestingly, *black* chromatin contains nearly two-thirds of all silent genes, most of them being tissue-specific genes, and appears to actively inhibit gene expression [5, 36]. How this repression is achieved is still unclear. Proteins that are now known to mark *black* chromatin are, notably, the linker histone H1, which has previously been linked to repression of transcription [36]. By cross-linking the entering and exiting DNAs of each nucleosome, H1 may indeed result in an effective shortening of linker DNAs [37], hence explain the stiffening and compaction of the nucleosome fiber. Moreover, these structural features sound reasonably related with gene silencing, hence giving further credibility to the hypothesis of H1 as the main actor in inactivation. In Fig. 3**D**, H1 proteins are sketched as green spheres on a two-angle model of an inactive, *black* Kuhn segment.

### 4.3 Looking for a molecular basis to explain the inferred energy parameters

Our approach provides the first *color*-specific inference of the interaction energy *ε* between chromatin Kuhn segments *in vivo*. In the original paper Ref. [2], a very high attractive interaction of 3.5 *k*_*B*_*T* is used in simulating the repressed domains. Because of this huge energy value, indeed, the globule conformations obtained by these authors are already close packed for *N* = 400 (Figure 4.c of Ref. [2]). Simulations are there only intended to reproduce the experimentally measured scaling behaviour, and give therefore only adimensional values for the radii of gyration. However, we can easily compare those results with our adimensional theoretical plot, as shown by the green dots in Fig. 1. It is clear from this comparison that such compact conformations could not fit *quantitatively* the experimental data. The globule volume can also be calculated for 2. 5 < *R*_*g*_ < 3 (approximately the limit point in Fig. 1) as 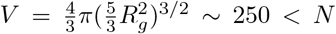: hence even denser than close packed.

Other estimations of chromatin interaction parameters have been obtained from the fit of Hi-C data [38, 39]. In a recent study, Falk *et al.* have determined the value of the interaction energy parameters in a copolymer model (A and B chromatin compartments) [40]. In order to recover the experimental phase separation between chromatin A and B, they found an interaction between B monomers of 0. 55 *k*_*B*_*T* and a much weaker interaction between A monomers. This is compatible with our results, assimilating the A compartment with active chromatin, and the B compartment with repressed ones.

Note that, since Fish hybridization implies DNA denaturation, a potential effect might be a partial chromatin decondensation. In this case, the effective interaction energy fitted by our procedure would be underestimated with respect to *in vivo* conditions. However, the FISH hybridization protocol adapted in [2] ensures minimal alteration of chromatin structure [41], as also indirectly indicated by the measurable folding of active, inactive and repressed domains.

It is therefore tempting to try to relate the different values of *ε* obtained for the three epigenetic states (Table 3) to different molecular interaction mechanisms. Caution is needed, since *ε* is an effective parameter accounting for the overall, mean interaction energy between two Kuhn segments. Simulations of nucleosome fibers with a fine-graining of 10 bp for DNA indicate that, on average, one should expect only one nucleosome-nucleosome contact *in trans* per Kuhn segment (Pascal Carrivain, personal communication). Assuming this, *ε* appears as a reasonable estimate for single *in trans* interaction, so that a direct comparison between the fitted values becomes possible. In the case of repressed domains, such interaction is known to be mediated by Polycomb proteins which are considered to stabilize condensed chromatin configurations by means of bridges, and we find, coherently, the largestinteraction energy *ε* ≃ 0. 4 *k*_*B*_*T*. It is however unclear which mechanism could explain the difference in interaction energy between active and inactive domains. As we will discuss in next section, several independent experiments provide a possible explanation, which does not involve protein-mediated interactions.

### 4.4 Comparison with nucleosome core particles solution experiments reveals criticality features and a key role for nucleosome-nucleosome interaction

The free energy expressed in terms of the renormalized density *t* = *ρ*^1/(*νd−*1)^ (with 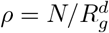, see Equation 4) comes from a *virial expansion* approach. This consists in assuming that interactions are dominated by two-body interactions, whereas many-body ones are rare, so that an expansion in terms of the (small) density parameter is suitable. In the case of polymers, at the coil-globule transition the second virial coefficient *a*_1_(*ε*) cancels out and changes its sign, reflecting a compensation between attractive and repulsive interactions, while *a*_2_(*ε*) remains positive [11]. Here we found that, if active domains are in the coil regime for all the observed lengths (with *ε* = 0. 10 *k*_*B*_*T*), inactive and repressed domains are, with *ε* = 0. 32 *k*_*B*_*T* and 0. 44 *k*_*B*_*T* respectively, in the crossover region for most of the observed lengths, due to finite-size effects. Hence, the second virial coefficient is close to zero for not-active domains, indicating a large degree of compensation between attraction and repulsion. Again, the question of the molecular basis of this behavior arises.

Worthwhilely, describing the system in terms of virial parameters allows us to compare our findings with completely independent experiments. In Refs. [8, 9], Livolant and co-workers experimentally characterized the interaction between isolated nucleosome core particles at different monovalent salt concentrations. Interestingly, the second virial coefficient steeply decreases to zero and presents a cusp in the salt range 75–210 mM, i.e. around physiological concentrations. Hence, the nucleosome architecture and biochemistry seem to have been selected so that repulsion and attraction between nucleosomes counterbalance in living organisms.

It is therefore tempting to link the coil-globule transition of chromosomes to the vanishing of the second virial coefficient of nucleosome-nucleosome interaction. This is also in line with quite recent measurements of chromosome dynamics in yeast, which has been modeled as a Rouse dynamics slowed down by nucleosome-nucleosome transient interactions [29]. Following this line and going into more detail, inactive (*black*) chromatin is very close to the Θ-point, indicating that nucleosome-nucleosome interactions might be dominant within inactive domains. For the active (*red*) chromatin, we propose that the lower interaction is linked to a lower interaction between nucleosomes, which is consistent with acetylation of histone tails in transcribing chromatin [42], thus reducing their charge, hence their ability to bridge other nucleosomes [43]. Structural changes of chromatin upon histone tail acetylation were indeed recently reported *in vitro* and *in vivo* [31, 44, 45]. For repressed (*blue*) chromatin, a larger value of *ε* points toward a stronger interaction, certainly mediated by proteins from the Polycomb family, in agreement with Ph-knockdown experiments [2]. The detailed modeling of the mechanistic effects involved remains elusive and clearly points to the need for molecular modeling of the Polycomb gene silencing complexes.

### 4.5 Subdomains and packing conditions

A last point to be discussed concerns the behavior of subdomains, *i.e.* internal regions of varying lengths within epigenetic domains. Boettiger *et al.* observe a plateau in the plot of 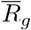 as a function of the genomic size for the subdomains of the two largest repressed domains. For these domains, 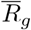 early saturates for subdomains that are about one fifth of the length of the parent domain. In all other cases (all active domains, all inactive domains and all other repressed domains), the plots of 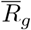 as a function of the subdomain length are the same as the plots of 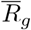 as a function of the domain length (see Figure 2b and Extended Data Figure 6b in Ref. [2]). This strongly supports the existence of a coil-globule transition: only the largest repressed domains are globular enough to exhibit a plateau in the subdomains plot. In all other cases, the conformations are coils either because the domains are above the Θ-point (active domains) or because they are too small to be globular (all *black* domains and all other *blue* domains).

The largest repressed domains behavior is described in Ref. [2] as *intermixing*. We note that this behavior is a well known characteristic of globular domains of polymers [46]: a chain inside a globule behaves like a random walk, until it hits the boundary of the globule. It then starts a new random walk inside the globule in such a way that, after several such collisions, the volume of the globule becomes filled with random walks that are uncorrelated with each other [47].

Boettiger *et al.* obtain an early plateau in their simulations that is comparable to the experimental plateau because they use a huge value of *ε* (larger than 1 *k*_*B*_*T*). Indeed, at *ε* of the order of 0.5 *k*_*B*_*T*, as we found for repressed domains, the plateau is reached at a much larger value of the subdomain length (about two thirds of the domain length, data not shown). We make the guess that the early saturation observed in experiments is probably due to the formation of loops mediated by Polycomb Response Elements (PRE) [48]. As these loops are rare, we do not expect them to have a significant impact on the results of our modeling [12]. This point deserves however further investigations and will be addressed in a future work.

## 5 CONCLUSION

Super-resolution imaging of chromosomal domains has opened a new era in modeling the 3D organization of nuclei. The whole distributions of structural properties of chromosomes, as the radius of gyration of epigenetic domains, are now available. We showed here how to use these distributions to get hitherto unavailable physical parameters of chromatin. In particular, we could get:

First, *color*-specific measures of the Kuhn length (in base pairs *and* in nanometers) of active, inactive and repressed domains respectively. Strikingly, these measures are on par with Hi-C data in mammals [34] as well as to most recent dynamic measurements in yeast [29]. The knowledge of both Kuhn lengths leads to the value of the compaction of the chromatin, *i.e.* the number of nucleosomes per 10 nm. This is a precious indication of the *conformational* state of the nucleosome fiber.

Second, we get the first measure of the interaction energy *ε* between Kuhn segments. It is very striking that, in all but two cases studied here (95%), the length of epigenetic domains remains small enough so that the domains are still in the coil region of the phase diagram. This suggests that one essential role of the coil-globule transition is to create *dense coils* which at the same time allow to “tidy up” a whole genome in the reduced volume of a cell nucleus while giving access in a reversible way to the transcription machinery. Importantly the high density of chromatin inside cell nuclei is not imposed by nuclear membrane confinement but by transient interactions.

It is often stressed that nucleosomes enable to reduce the length of a chromosome by a factor of ten. Our results point to a new role of the nucleosome: *nucleosome-nucleosome interaction is by itself strong enough to induce chromosome folding at the level of epigenetic domains*, hence to drive chromatin organization. We find indeed that the specific value of the interaction energy between nucleosomes may allow by itself the existence of a coil-globule transition in the neighborhood of typical physiological conditions, in particular for inactive domains where no clear evidence for protein-mediated interactions directly affecting the folding state has been reported until now [36]. Interaction between nucleosomes have been directly observed in recent cryo-EM experiments [49], pointing out in particular a central role of H3 and H4 histone tails. In mammals, perturbation experiments already indicated that chromatin domains are organized by a combination of cohesin and nucleosome-nucleosome interactions, the latter being in particular tuned by histone tail acetylation [31]. The key role of histone tail flexibility on chromatin compaction has also been shown by computational studies [43]. We therefore speculate that histone tail sequences have been selected by evolution for nucleosome-nucleosome attraction to be close to repulsion in physiological conditions. This may explain why histone tail sequences are so well conserved among eukaryotes, although they are intrinsically disordered protein domains [50]. To further consolidate this assumption on nucleosome-nucleosome interaction, it would be highly desirable to manipulate in vivo histone-tail modifications or histone variants in a similar way to the one initiated by Boettiger *et al.* for Polycomb repressed domains. Interestingly Gibson *et al.* just showed in vitro that nucleosome arrays undergo liquid-liquid phase separation (LLPS) which is inhibited by histone acetylation and promoted by linker histone H1 [51]. Super-resolution microscopy [2, 3, 52] combined with the methodology presented in this paper now allows to design new experiments to investigate the effect of such molecular modifications on the 3D organization of chromatin sub-compartments.

## Acknowledgements

We are very grateful to Alistair Boettiger, Xiaowei Zhuang and co-workers for kindly providing us with their experimental data and for interesting discussions. We thank Giacomo Cavalli and Marcelo Nollmann for helping us to understand some aspects of the experimental approach. Very relevant and constructive Referees’ comments are also sincerely acknowledged. This work is dedicated to the memory of our colleague and friend Alain Arneodo. His enthusiasm and passion for knowledge and interdisciplinarity have accompanied our work for many years and will always remain a guide for us.

## Footnotes

[1] Simulations are performed on a cubic lattice. The conformations of a polymer of *N* monomers are sampled thanks to the Metropolis algorithm with reptation moves [19]. The overall conformation energy is obtained by weighting the number of contacts by an energy cost per contact *−ε*. Incidentally, theoretical expression and on-lattice simulations are in excellent agreement [17].

