## Supplemental figures for "Polymer coil-globule phase transition is a universal folding principle of Drosophila epigenetic domains"

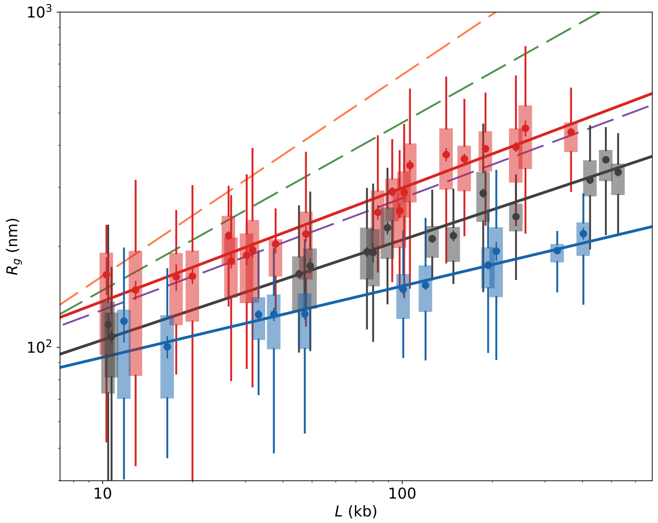


**(a)**

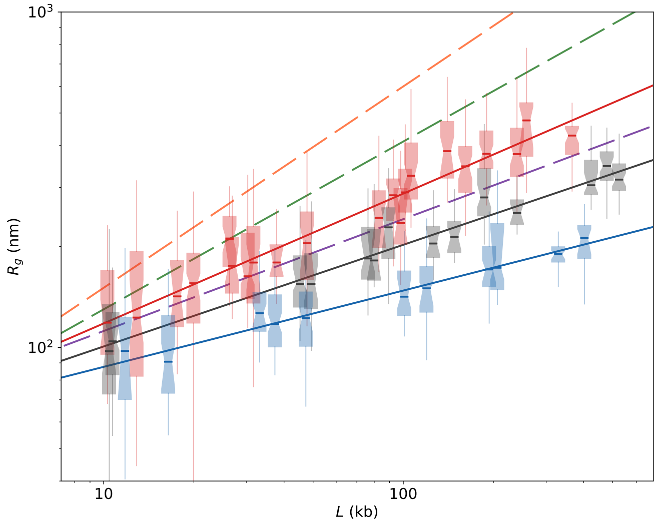


**(b)**

**Figure S1** Radii of gyration $\overline{R}_g$ calculated from the dataset of Boettiger et al 2016 for the three sets of domains: active (red), inactive (black) and repressed (blue). **(a)** Data are represented as boxplots, showing the interquartile range (IQR : 25 to 75 percentile), with whiskers adding and subtracting additional 1.5 IQR. Points indicate the mean, errorbars are the standard error of the mean (i.e. the sample standard deviation over the square root of the sample size). Solid lines are power laws ($A\,L^b$) fitted to the series of mean values, weighted by standard errors of the mean. Corresponding Flory exponents $\nu$ are $0.34$ (active), $0.30$ (inactive), $0.19$ (repressed). **(b)** Medians from the same datasets are indicated by the horizontal segments. Notches display the confidence interval around the median based on the median $+/-$ 1.57 IQR over the square root of the sample size. Solid lines are power laws ($A\,L^b$) fitted to the series of median values. Corresponding Flory exponents $\nu$ are $0.36$ (active), $0.31$ (inactive), $0.22$ (repressed). **(a, b)** Dashed lines, shown for reference, represent the typical scaling laws for random walk polymers ($\nu = 0.5$, green), self-avoiding walk polymers ($\nu = 3/5$, orange) and globule polymers ($\nu = 1/3$, blue).

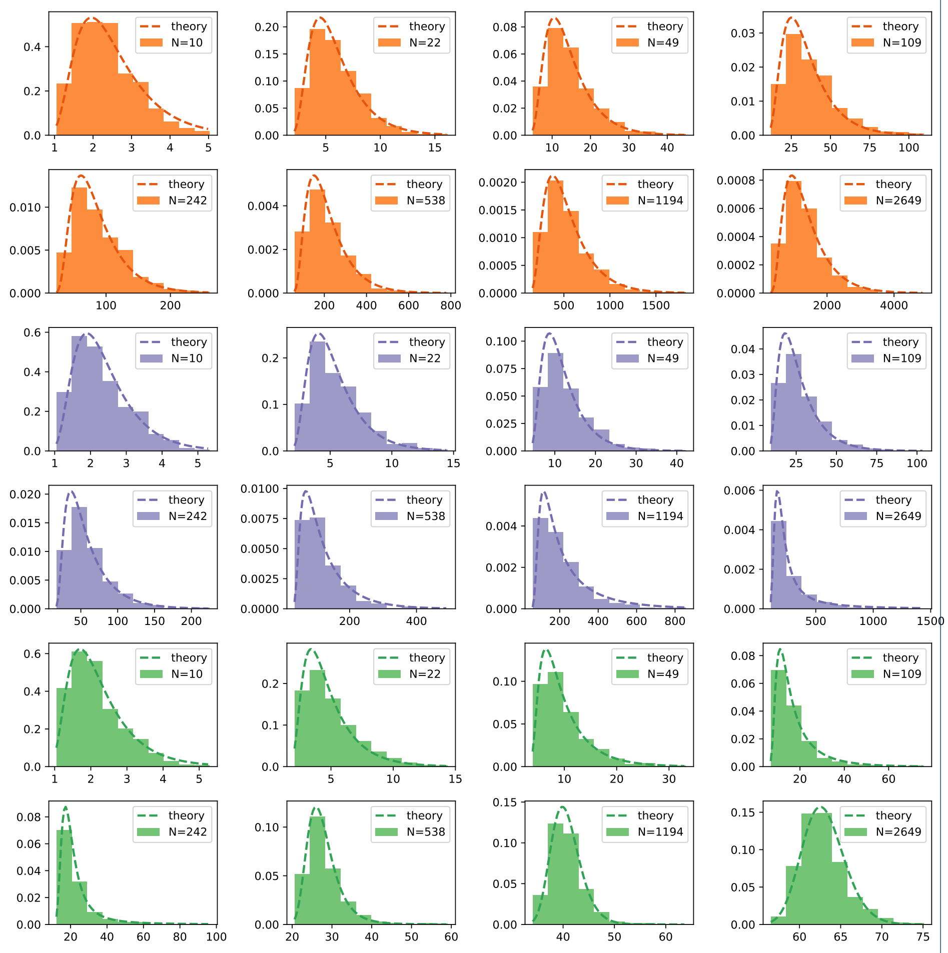


**Figure S2** Selected histograms from the on-lattice simulations for different polymer lengths $N$ from 10 to 2649, obtained with interaction energies $\varepsilon = 0.20$ (orange), $\varepsilon = 0.31$ (purple) and $\varepsilon = 0.44$ (green). Dashed lines correspond to the theoretical model of the free energy $\mathcal{F}_N(t|\varepsilon) = a_1(\varepsilon)Nt + a_2(\varepsilon)Nt^2 + a_3(\varepsilon)(Nt)^{-2/3} + a_4(\varepsilon)(Nt^2)^{2/3} + 1.13\ln Nt$.

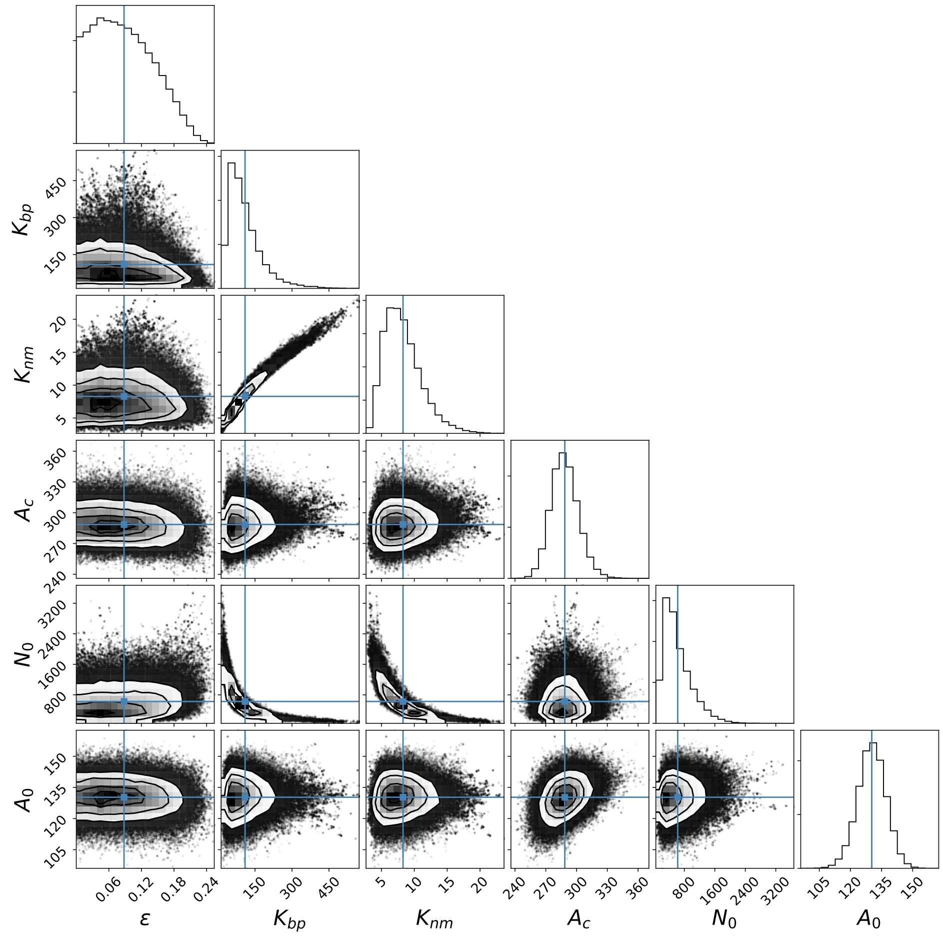


**Figure S3** Parameter distribution inferred from data by Bayesian analysis for active (red) domain measurements. On the diagonal: parameter distributions, marginalized on all other parameters. Off diagonal: distributions marginalized on each parameter couple (see legends). In all graphs, light blue lines indicate the position of the mean value of the corresponding parameters, used as best parameter estimates in Table 3.

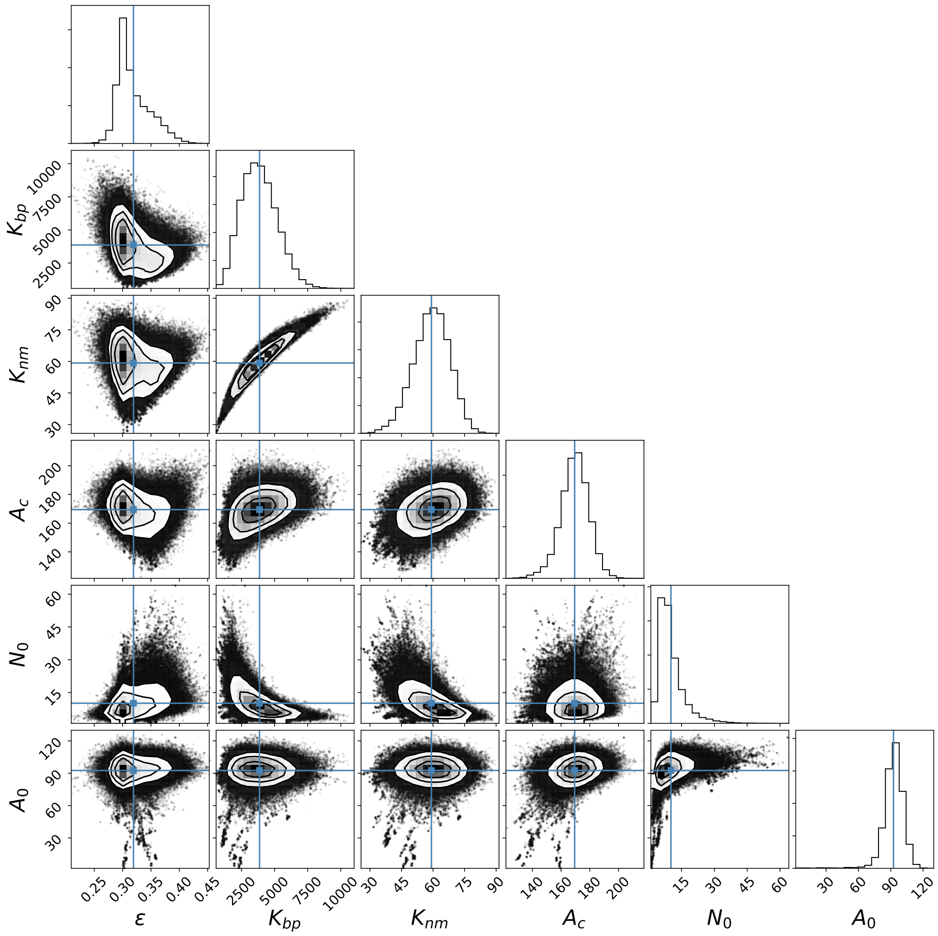


**Figure S4** Parameter distribution inferred from data by Bayesian analysis for inactive (black) domain measurements. On the diagonal: parameter distributions, marginalized on all other parameters. Off diagonal: distributions marginalized on each parameter couple (see legends). In all graphs, light blue lines indicate the position of the mean value of the corresponding parameters, used as best parameter estimates in Table 3.

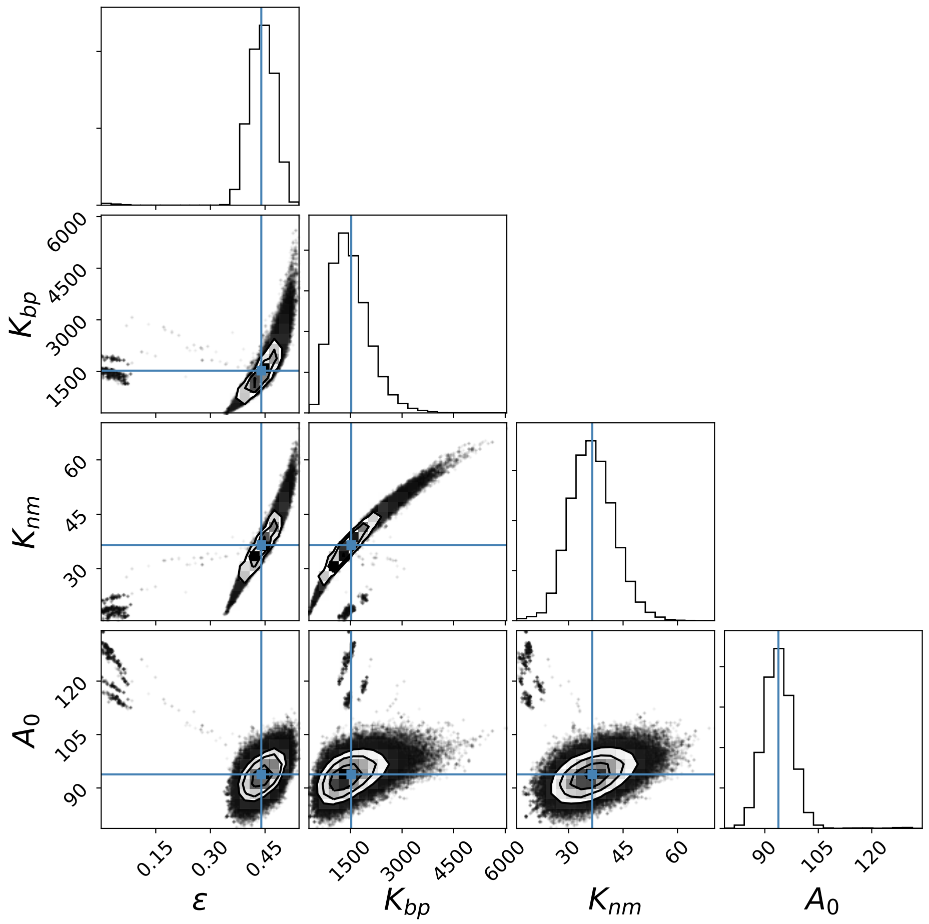


**Figure S5** Parameter distribution inferred from data by Bayesian analysis for repressed (blue) domain measurements. On the diagonal: parameter distributions, marginalized on all other parameters. Off diagonal: distributions marginalized on each parameter couple (see legends). In all graphs, light blue lines indicate the position of the mean value of the corresponding parameters, used as best parameter estimates in Table 3.

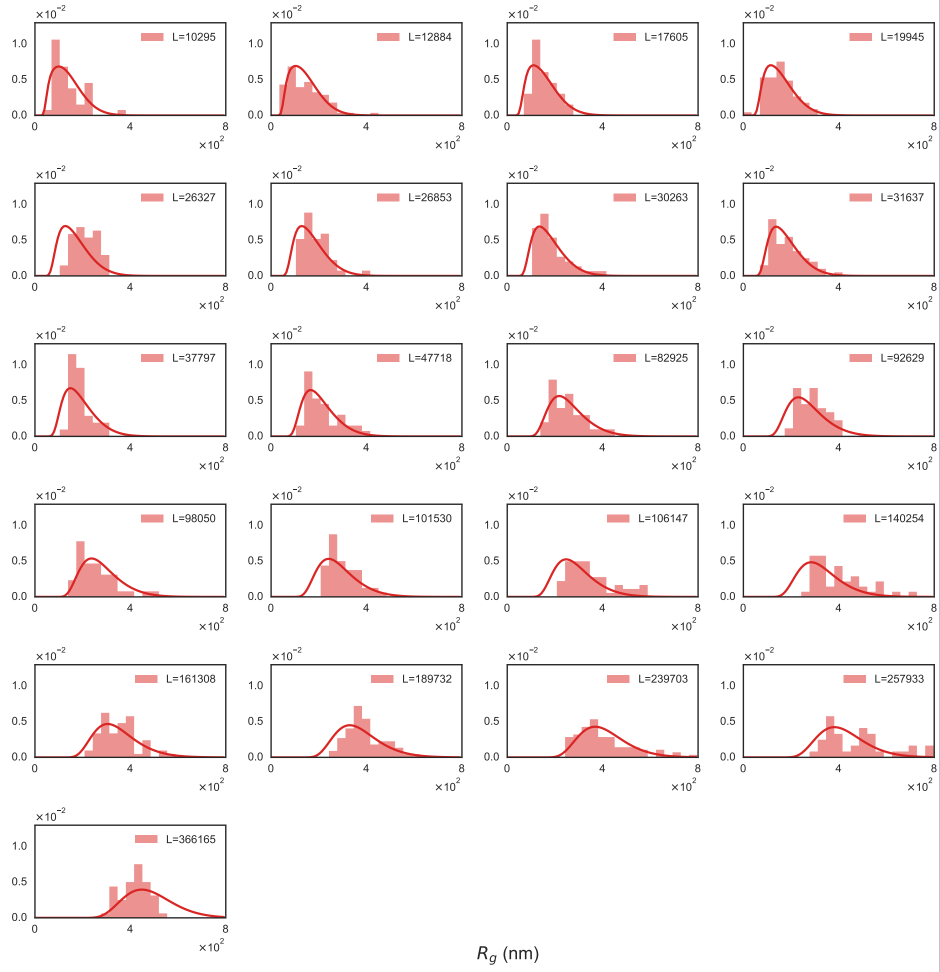


**Figure S6** The ensemble of data histograms with the corresponding theoretical distributions obtained using the parameters obtained from the fit for active domains.

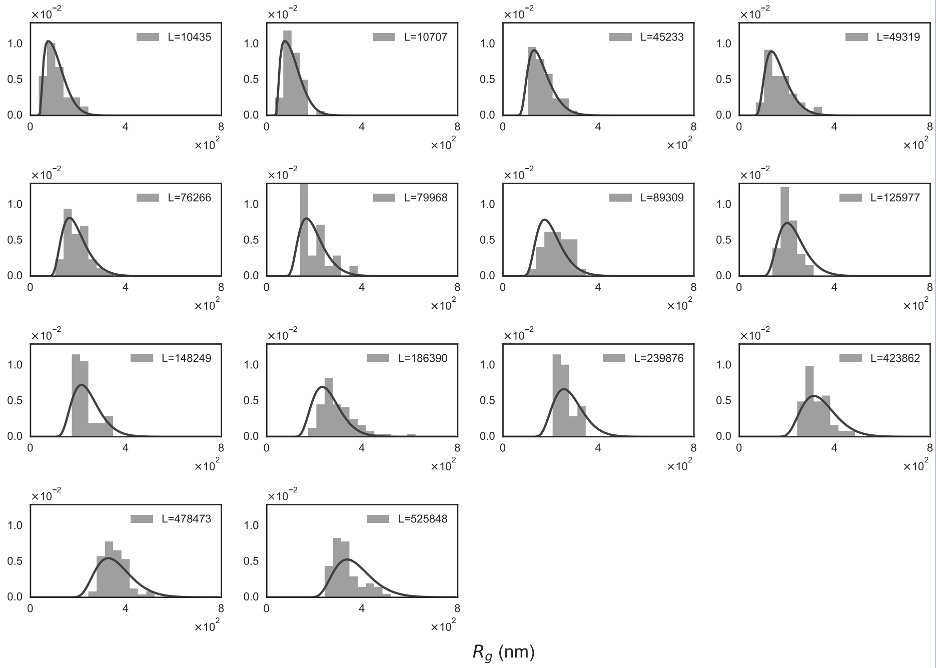


**Figure S7** The ensemble of data histograms with the corresponding theoretical distributions obtained using the parameters obtained from the fit for inactive domains.

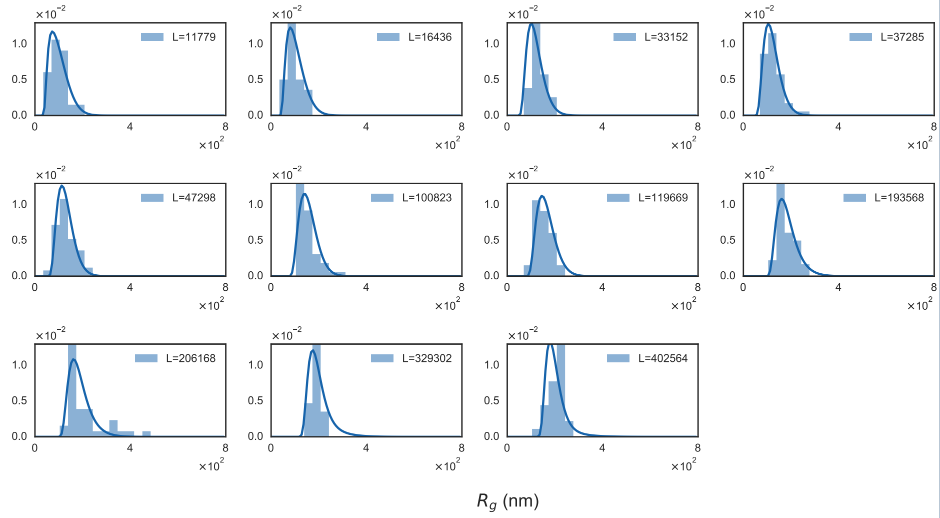


**Figure S8** The ensemble of data histograms with the corresponding theoretical distributions obtained using the parameters obtained from the fit for repressed domains.

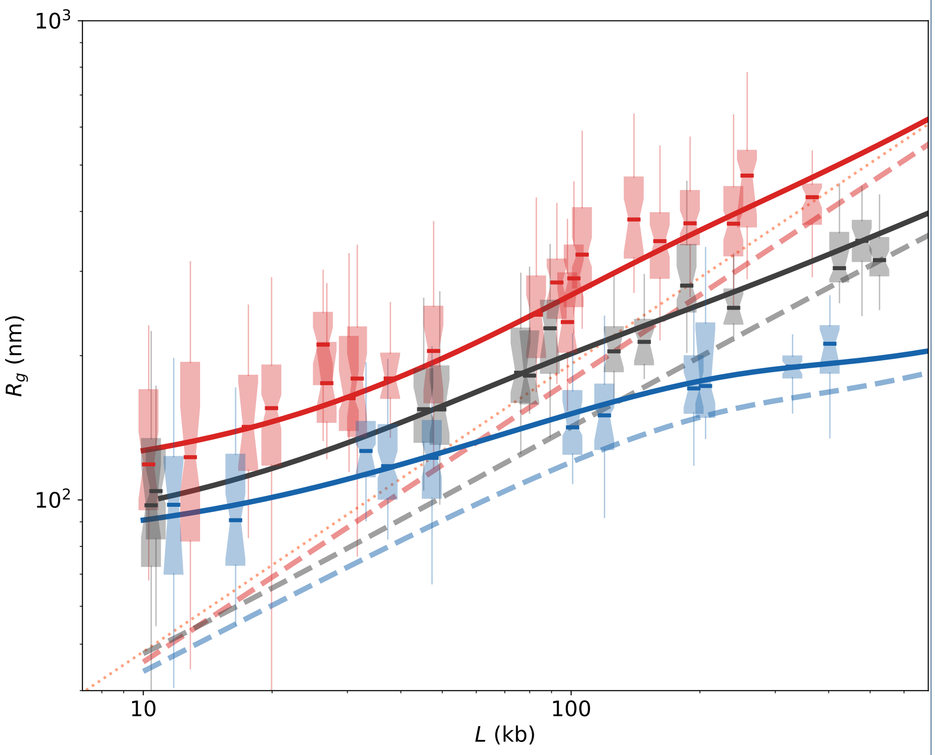


**Figure S9** *Medians* of $R_g$ as a function of the domain length $L$ calculated from the analytical model with the parameter sets of Table 3: active (red line), inactive (black line), repressed (blue line). Boxplots (same colors respectively) correspond to the experimental data from Boettiger et al 2016, as in Fig. S1. Medians are indicated by the horizontal segments. Dashed lines are obtained from previous fitting curves by deconvolution, hence correspond to the behavior expected in a typical haploid system. The orange dotted line represents the coil ($\nu = 3/5$) typical scaling law. Complementary information with respect to Fig. 3 A.
